# Rapid pacing by circulating traveling waves improves maturation of hiPSC-derived cardiomyocytes in self-organized tissue ring

**DOI:** 10.1101/717108

**Authors:** Junjun Li, Lu Zhang, Leqian Yu, Itsunari Minami, Marcel Hörning, Ji Dong, Jing Qiao, Nanae Fujimoto, Yuji Shiba, Yang Zhao, Fuchou Tang, Shigeru Miyagawa, Yong Chen, Yoshiki Sawa, Chao Tang, Li Liu

## Abstract

Directed differentiation methods allow acquisition of high-purity cardiomyocytes (CMs) differentiated from human induced pluripotent stem cells (hiPSCs); however, their immaturity characteristic limits their application for drug screening and regenerative therapy. The rapid electrical pacing of cardiomyocytes have been used for efficiently promoting the maturation of cardiomyocytes, here we describe a simple device in modified culture plate on which hiPSC-derived CMs (hiPSC-CMs) can form three-dimensional self-organized tissue rings (SOTRs). Using calcium imaging, we show that within the ring, traveling waves (TWs) of action potential spontaneously originated and ran robustly at a frequency up to 4 Hz. After 2 weeks, SOTRs with TW training showed matured features including structural organization, increased cardiac-specific gene expression, enhanced Ca^2+^-handling properties, an increased oxygen-consumption rate, and enhanced contractile force. We subsequently used a mathematical model to interpret the origination, propagation, and long-term behavior of the TWs within the SOTRs. This new idea for spontaneous hiPSC-CM maturation also has potential for pacing the electrical excitable cells such as neuron and retina cells for various applications.

## Introduction

Although human induced pluripotent stem cell-derived cardiomyocytes (hiPSC-CMs) have been proposed as an abundant resource for tissue engineering, drug screening, and regenerative-medicine applications^1^, they exhibit characteristics different from adult human CMs, including immature sarcomere structure and morphology, a fetus-like gene-expression profile, and inadequate Ca^2+^-handling properties^2–4^. The immature nature of hiPSC-CMs potentially hinders their reflection of adult heart physiology for disease modeling and drug assessment.

To achieve higher degrees of CM maturation and function, a variety of methods have been developed, including dynamic culture^5^, three-dimensional (3D) engineered heart tissue^6,7^, 3D printing^8^, addition of factors^9^, extracellular matrix^9,10^, and external stimulation. Electrical stimulation, especially high-frequency pacing, has long been suggested as an effective method for maturing muscle cells^11–14^, which upregulates cardiac markers^12,13,15^, enhances Ca^2+^-handling properties^3^, and promotes CM alignment^16^. During this process, appropriate stimulation protocols are required to minimize possible tissue damage^17^. Additionally, it remains challenging to rapidly pace CM tissues (≥2 Hz) over long time^18^ due to possible side effects, such as heavy metal poisoning, electrolysis, pH shift, and the generation of reactive oxygen species (ROS)^19,20^. Moreover, upscaling for mass stimulation is either difficult or requires high level of power consumption^18^. As an important complement, mechanical stimulation can promote CM maturation, with cyclic mechanical stress applied by external stretching devices capable of forcing muscle-cell assembly into structurally and functionally aligned 3D syncytium^21^, thereby affecting their gene expression^22–24^ and Ca^2+^ cycling^25^.

On another hand, spiral waves (traveling waves) within cardiac tissue have long been investigated as a model for arrhythmia studies^26–29^, and the rapid beating caused by in-vivo spiral wave could lead to dysfunction of heart and even death of a patient. However, when used in in-vitro condition, the rapid pacing by spiral wave, similar to rapid electrical stimulation, could be beneficial for cardiomyocytes maturation.

In this study, we created a platform capable of promoting rapid formation of hiPSC-CMs into 3D self-organized tissue rings (SOTRs), where propagation of an action potential in the form of traveling waves (TWs) can spontaneously originate and travel around the close-looped circuit, thereby making the CMs beat at a high frequency (~2–4 Hz) continuously and robustly up to more than 89 days without any external stimulation. Additionally, we found that the beating frequency and the TW speed could be adjusted by changing the diameter of the ring. Furthermore, we constructed a mathematical model in order to elucidate the origination, propagation, and long-term behavior of the TWs, with the model ultimately agreeing well with the experimental data. After 2 weeks of training using the TWs, the SOTRs demonstrated improved structural organization, upregulated cardiac-specific gene expression, enhanced Ca^2+^-handling properties, an increased oxygen-consumption rate (OCR), and enhanced contractile force.

Although the maturation level of CMs trained with TWs remains lower than that acquired using state-of-art methods^13^, the maturation process demonstrated in the present study was spontaneous and required no elaborate experimental setup and/or external stimulation. Moreover, the mass production of matured CMs would be possible by simply modifying conventional Petri dishes. Our results demonstrated a novel approach for promoting spontaneous CM maturation and could serve as an economical and practical system for future production of matured hiPSC-CMs.

## Results

### Self-organization of a hiPSC-CM ring

We created 3D SOTRs by plating hiPSC-CMs in a culture dish with a pillar in the center, around which the CMs aggregated and formed a thick tissue ring within 2 days (Fig. 1a–c and Supplementary Fig. 1). As indicated by the genetically encoded calcium indicator GCaMP, we found looped activation propagations within the SOTRs (i.e., TWs; Supplementary Video 1), with from zero to three TWs present in one SOTR (Fig. 1d and e; and Supplementary Video 2).

**Figure 1.**
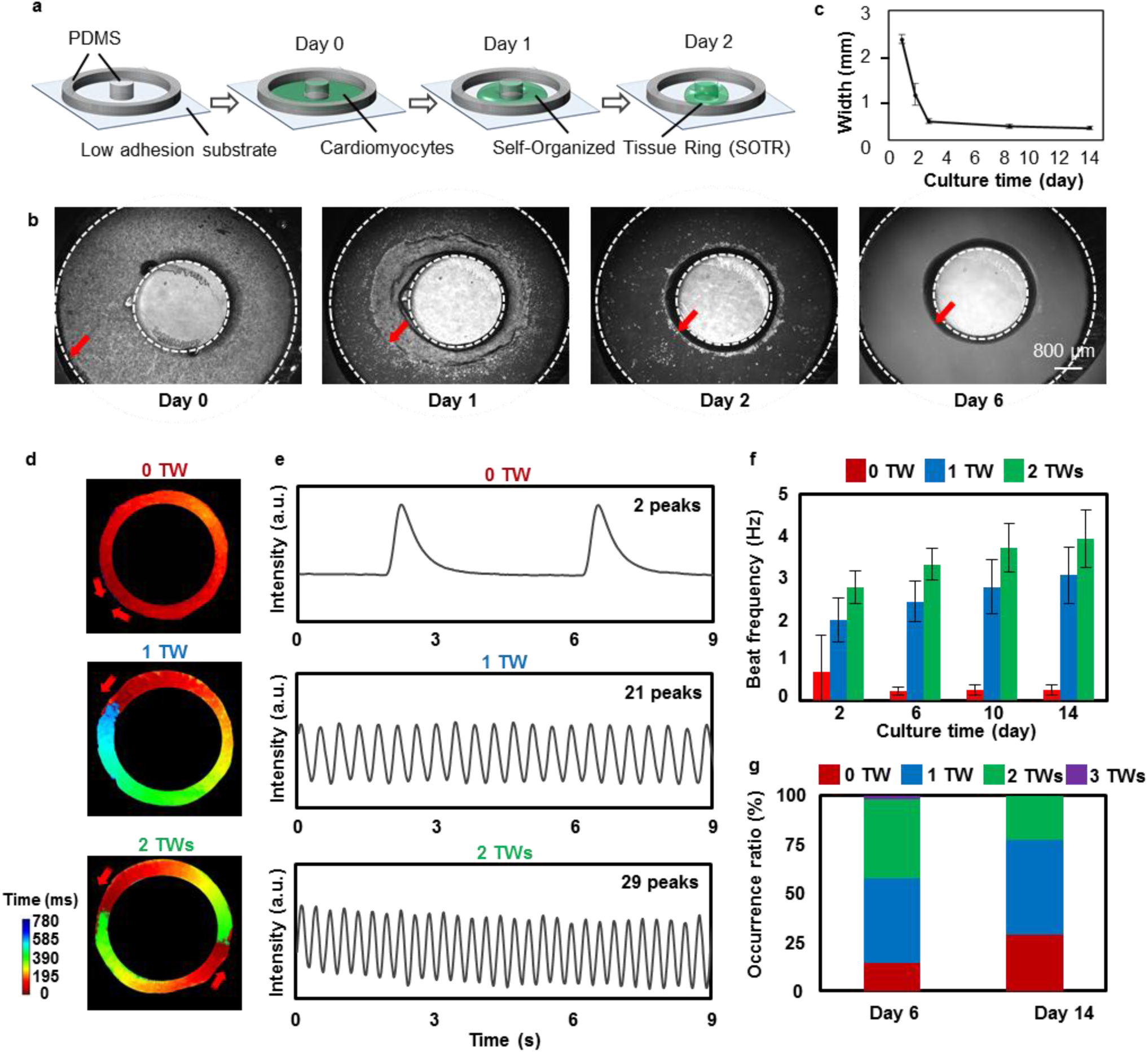
TWs promoted rapid beating of cells in SOTRs. (a) Schematic describing SOTR formation. (b) Bright-field images of hiPSC-CMs in the template. The red arrows indicate the edge of cardiac tissues in the template, and the dashed lines indicated the PDMS block and pillar boundaries, respectively. Pillar diameter = 3mm. (c) Quantification of the width of the SOTRs on the indicated culture day (mean ± s.d.; *n* = 4). (d) Activation map of GCaMP3-positive SOTRs with zero, one, or two TWs. The red arrows indicate the propagation direction of the action potential. (e) GCaMP3-fluorescence signal at a fixed position on the ring of SOTRs with zero, one, or two TWs on day 6. (f) Beat rates of SOTRs at differentculture times (mean ± s.d.; 2 TWs: *n* = 10; 1 TW: *n* = 12; 0 TWs: *n* = 8). (g) The percentage of occurrence for different numbers of TWs on days 6 (n = 204) and 14 (n = 186), respectively.

A trace recording (Fig. 1d and e) indicated no long rest periods between successive TWs, resulting in higher beating rates in SOTRs capable of sustaining a higher number of TWs (Fig. 1e and f). The spontaneous beating rate of SOTRs with zero TWs was 0.21 ± 0.10 Hz on day 6 and remained stable for 2 weeks, whereas the rates of SOTRs with one or two TWs on day 6 were much higher (2.40 ± 0.49 Hz and 3.30 ± 0.39 Hz, respectively). Notably, the wave speed in the TW groups was much lower (~2 cm/s) than that of the spontaneous beating group (0 TW; ~9 cm/s). It is possible that the slower speed together with the shortened pacing interval and refractory period might be caused by the higher beating frequency in the TW groups relative to that in the zero TW group (Supplementary Fig. 2), which agrees with previous reports associated with excitable media^30–34^.

The beating rates of the one and two TW groups increased slightly over 2 weeks of culturing, reaching 3.03 ± 0.68 Hz and 3.92 ± 0.69 Hz on day 14, respectively. After a 6-day culture, 83.8% of 204 SOTRs had one or two TWs, 14.2% had zero TWs, and 2% had three TWs; however, these percentages changed after a 2-week culture to 28.5% with zero TWs, 48.9% with one TW, 22.6% with two TWs, and no SOTRs with three TWs (Fig. 2d). Noise and disturbances, such as those during medium changing, might have provoked changes in and/or disappearance of TWs. In the long-term culture, we found that TWs could be maintained in SOTRs for >89 days (Supplementary Fig. 3).

**Figure 2.**
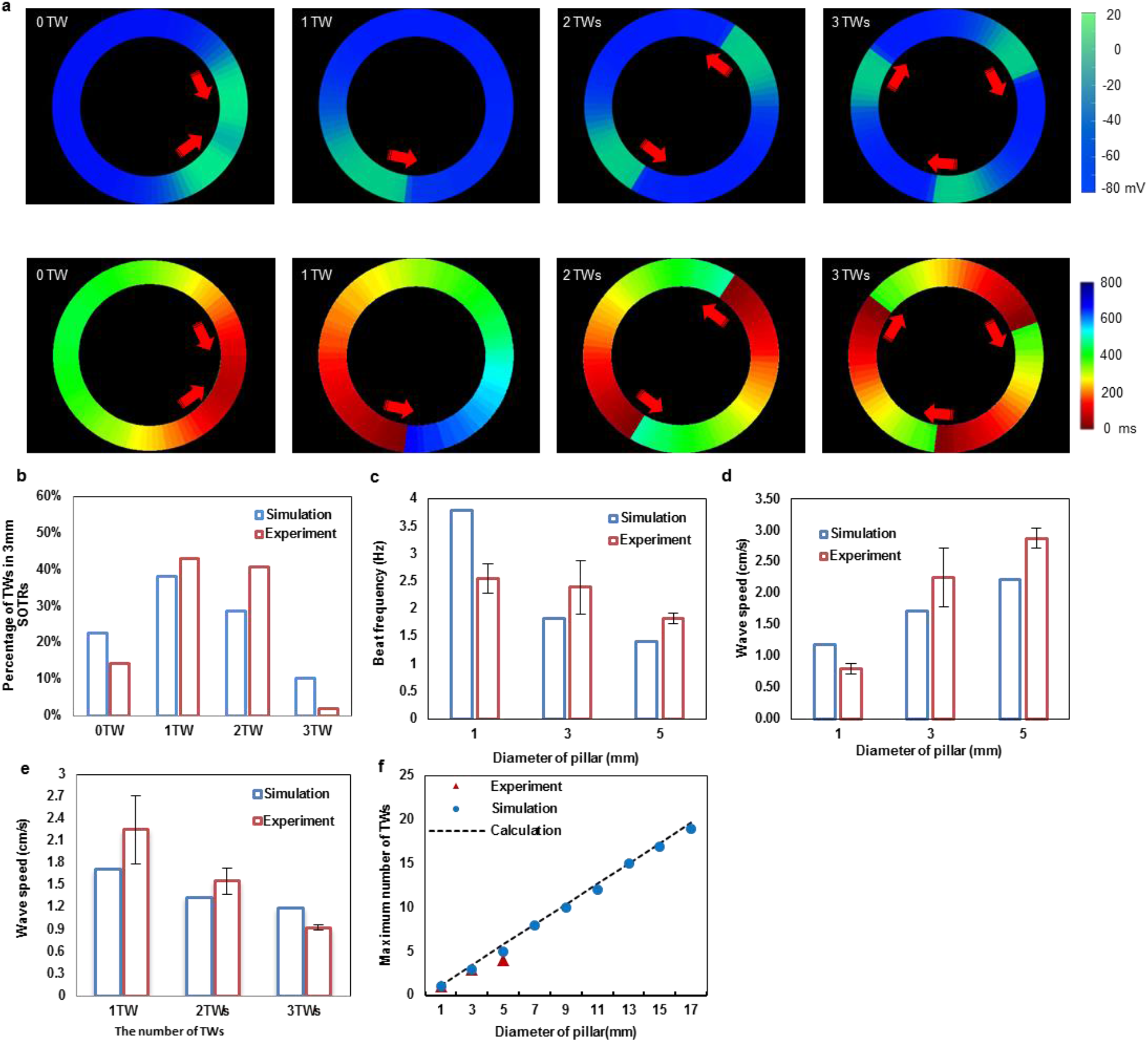
Accurate reproduction of TW features in SOTRs using a mathematical model. (a) Examples of simulation results of the model (upper panel) and the activation map (lower panel). For the group with zero TWs, an action potential initiates from the left position of the ring and propagates in the opposite directions along the ring, after which the two waves meet and annihilate on another. The process will starts again with new waves initiating at the same position periodically. For groups with one, two, or three TWs, stable TWs travel around the ring. Membrane potential and activation time are color coded. (b) The percentage of stable TWs in SOTRs at day 6 (diameter = 3 mm; *n* = 60 in the simulation and *n* = 204 in the experiment). (c) Beat rate (Hz) of CMs in SOTRs with one TW at day 6 for different pillar diameters [experiment: 1-mm SOTR (n = 4); 3-mm SOTR (n = 12); 5-mm SOTR (n = 10)]. (d) Similar information shown in (c) but with TW speed. (e) Speeds of one, two, or three TWs in SOTRs with a pillar diameter of 3 mm [experiment: mean ± s.d.; 3 TWs (n = 4); 2 TWs (n = 10); 1 TW (n = 12)]. (f) The maximum number of TWs in SOTRs with different pillar diameters.

To better understand the nature of these TWs according to their organization, propagation, and long-time behavior, we constructed a mathematical model (Supplementary Information and Supplementary Fig. 4) comprising a ring of cells, each of which can beat spontaneously with an intrinsic frequency. The total number of the cells in each ring in the model is proportional to the diameter of the ring. The coupling between neighboring cells through gap junctions^35,36^ gradually increases with time in order to simulate the selforganization and formation processes of the SOTR. Initially, all cells were independently set to a random phase of beating, with all cells beating independently. As the gap junctions form and strengthen, the beating of one cell harbors an increasing probability of triggering beating in its neighboring cell, thereby forming a propagating wave. Initially, these waves are unsynchronized and short-lived, as they initiate, disappear, and sometimes meet and collide (Supplementary Video 3). Over time, a SOTR is left with one dominant propagating-wave mode comprising zero or more TWs (Fig. 2a and Supplementary Video 3), with this process similar to that observed in our experimental findings (Supplementary Video 4). Notably, SOTRs with one or two TWs accounted for most of the simulation samples, a trend also observed in our experimental results (Fig. 2b).

We then investigated how TW features and properties were affected by characteristics, such as ring diameter, in the mathematical model and experiments. The beating frequency of cells in SOTRs decreased with increasing ring diameter, as shown with one TW in Fig. 2c. We noted that a constant TW speed implied a linear decrease in beating frequency with ring diameter (which is proportional to the ring perimeter), and that the greater the perimeter, the more time the wave would spend to travel around it. However, the data in Fig. 2c show a slower-than-linear decrease; therefore, we measured the wave speed, finding that it increased along with ring diameter in both the simulation and experiment (Fig. 2d). As discussed in the Supplementary Information, this was due to the ability of a single hiPSC-CM to spontaneously beat. Cells in larger rings would wait longer for the next TW to arrive, thereby making them easier to be activated by the wave front and resulting in a faster wave speed. This insight allowed us to predict the speed of two and three TWs. As shown in Fig. 2e, the prediction agreed well with our experimental results. Additionally, the mathematical model showed that the maximum number of TWs that a SOTR could sustain increased along with ring diameter, which was confirmed experimentally (Fig. 2f). Here, the mathematical model predicted that SOTRs with a diameter of 17 mm could contain as many as 19 TWs. Experimentally, we found that SOTRs with a 3-mm pillar were optimal based on their highest occurrence of TWs (~90% at ≥1 TW) (Supplementary Fig. 5). Moreover, the beating frequency of one TW was only slightly slower than that in SOTRs with a 1-mm pillar; therefore, SOTRs with a 3-mm pillar were chosen for subsequent investigations.

### TWs promote genetic and structural maturation of hiPSC-CMs

The human fetal heart rate varies, although it generally stabilizes at ~3 Hz, whereas the adult human heart rate is ~1 Hz^37^. Electrical stimulation has been used to mature CMs by pacing their beating at a certain higher-than-normal frequency relative to a normal human rate^3^; however, continuous and high electrical stimulation can cause cell damage. Here, TWs within the SOTRs were able to make the CMs beat at various frequencies in the absence of external stimulation.

We used RNA sequencing to compare gene-expression profiles among different groups. Principal component analysis (PCA; Fig. 3a) and hierarchical clustering of Spearman correlation results (Fig. 3b) showed closer correlations between hiPSC-CMs receiving training by TWs as compared with those without TW training. Additionally, we found different gene-expression patterns among zero, one, and two TW groups (Fig. 3c). Functional annotation according to Gene Ontology (GO) analysis revealed that upregulated genes (adjusted P < 0.05; fold change > 1.5) in SOTRs with two TWs relative to those with no TWs were related to the response to unfolded protein and a number of maturation-related terms.

**Figure 3.**
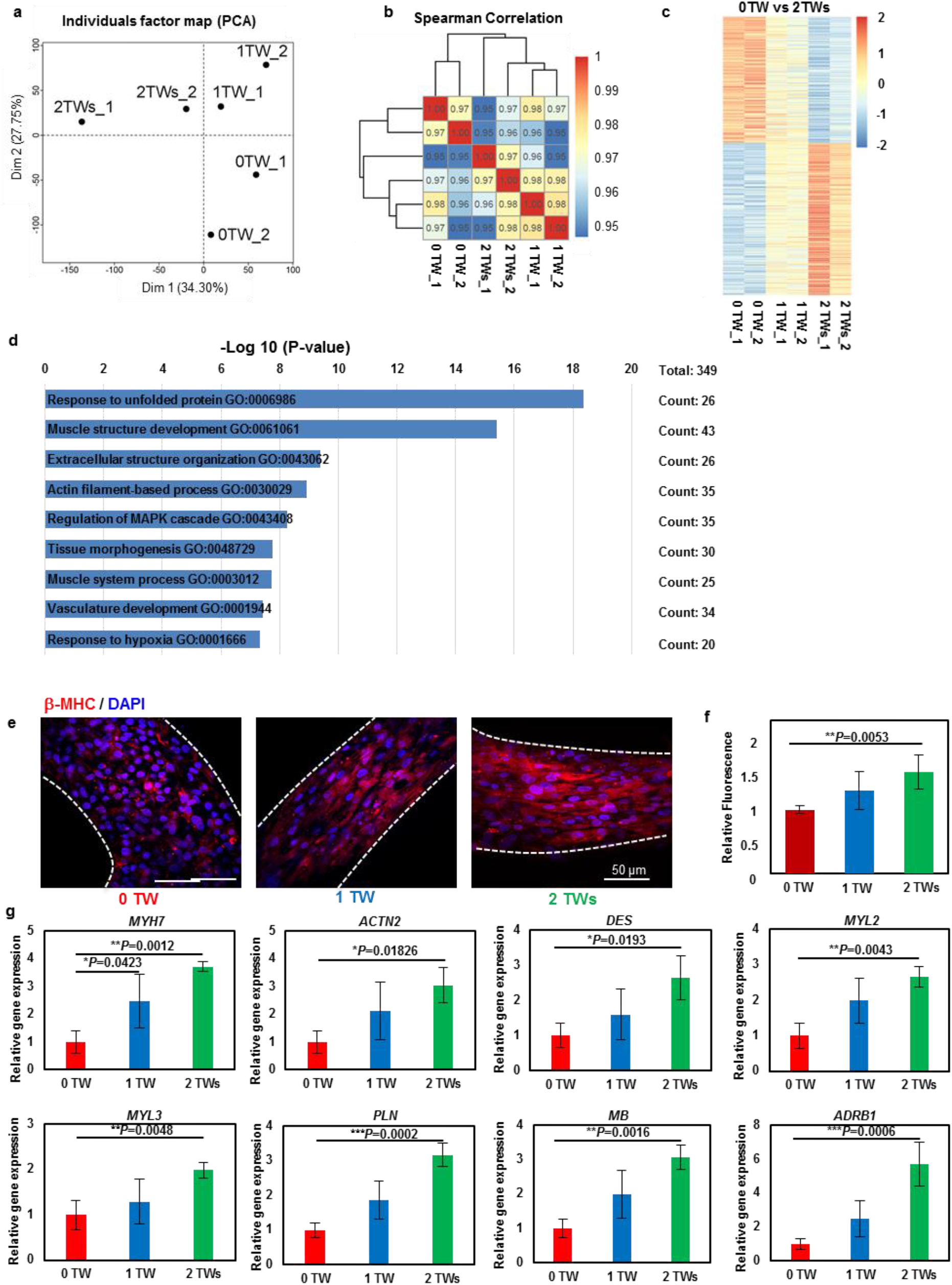
TWs upregulate cardiac-specific gene expression. (a) PCA of SOTRs with or without TWs based on RNA-sequencing data. (b) Heatmap showing hierarchical clustering of the correlation matrix resulting from comparison of expression values for each sample. Correlations calculated using Spearman correlation. (c) Heatmap showing the relative expression levels (z-score) of 349 genes from SOTRs with TWs (1 and 2 TWs) as compared with SOTRs with zero TWs. (d) GO categories significantly enriched for genes upregulated in SOTRs with two TWs as compared with SOTRs with zero TWs. Genes with fold changes > 1.5 and a P < 0.05 were analyzed. Enriched terms are listed. (e) Representative confocal images of SOTRs with zero, one, or two TWs on day 14. CMs were stained with anti-β-MHC (red) and DAPI (blue). (f) Quantification of β-MHC levels in SOTRs on day 14. Total integrated fluorescence of β-MHC was normalized to that of SOTRs with zero TWs (mean ± s.d.; *n* = 4). \*\**P* < 0.01. (g) The expression of cardiac-specific genes in SOTRs 14 days after cell seeding (mean ± s.d.; *n* = 4). \**P* < 0.05; \*\**P* < 0.01; \*\*\**P* < 0.001.

Most of the GO-enriched terms were associated with cardiac development, including muscle-structure development, extracellular-structure organization, actin-filament-based processes, tissue morphogenesis, and muscle-system processes. The response to unfolded protein is a coping response to mitigate or eliminate endoplasmic reticulum (ER) stress caused by hypoxia^38^, which could result from the high-frequency beating of the SOTRs with TWs. To verify these findings, we performed immunostaining and quantitative polymerase chain reaction (qPCR) analysis. Immunostaining for cardiac-specific markers clearly revealed higher expression of β-myosin heavy chain (β-MHC), a cardiac maturity marker correlated with contractile velocity, within SOTRs with one and two TWs as compared with levels observed in the zero TW group (Fig. 3e and f). Moreover, analysis of mRNA expression revealed a number of upregulated genes, including *MYH7* (encoding β-MHC) and genes involved in sarcomere structures *(ACTN2* and *DES*), ventricular structures (*MYL2, MYL3*), ER-Ca^2+^ function *(PLN), myoglobin (MB),* and *(β1-adrenoceptor (ADRB1).* SOTRs with TWs had significantly higher-fold upregulated genes as compared with those with zero TWs (Fig. 3e-g) due to higher beating rates during culture, which agreed well with previous reports utilizing electrical stimulation at different frequencies^3,15^.

Hematoxylin and eosin (HE) staining showed that CMs with TWs densely packed with each other to a greater degree than those without TWs (Supplementary Fig. 6). Moreover, SOTRs with TWs demonstrated densely packed cardiac myofilaments along the ring orientation, which was also along the path of the TWs, whereas CMs in SOTRs without TWs (0 TW) were randomly oriented and poorly organized (Fig. 4a and b). Additionally, after a 14-day culture, CMs in SOTRs with TWs strongly expressed the α-actinin as compared with those in SOTRs without TWs (Fig. 4c), in agreement to gene expression *(ACTN2,* Fig. 3g). Importantly, CMs in SOTRs with TWs exhibited significantly longer sarcomere length than those without TWs (1 TW: 1.71 ± 0.08 μm; 2 TWs: 1.83 ± 0.10 μm; and 0 TW: 1.49 ± 0.08 μm) (Fig. 4c and d), which is closer to the sarcomere length in human adult CMs^39^. Furthermore, electron microscopy indicated that cells in TW groups exhibited larger sarcomeric bundles, well-defined Z disks, A-bands, and myofibrils (Fig. 4e). These data suggested that CM maturation was improved within SOTRs with TWs relative to those with zero TWs. Our findings confirmed the genetic and structural maturation of SOTRs with TWs, and that faster beating rates during short-term culture could lead to higher levels of maturation.

**Figure 4.**
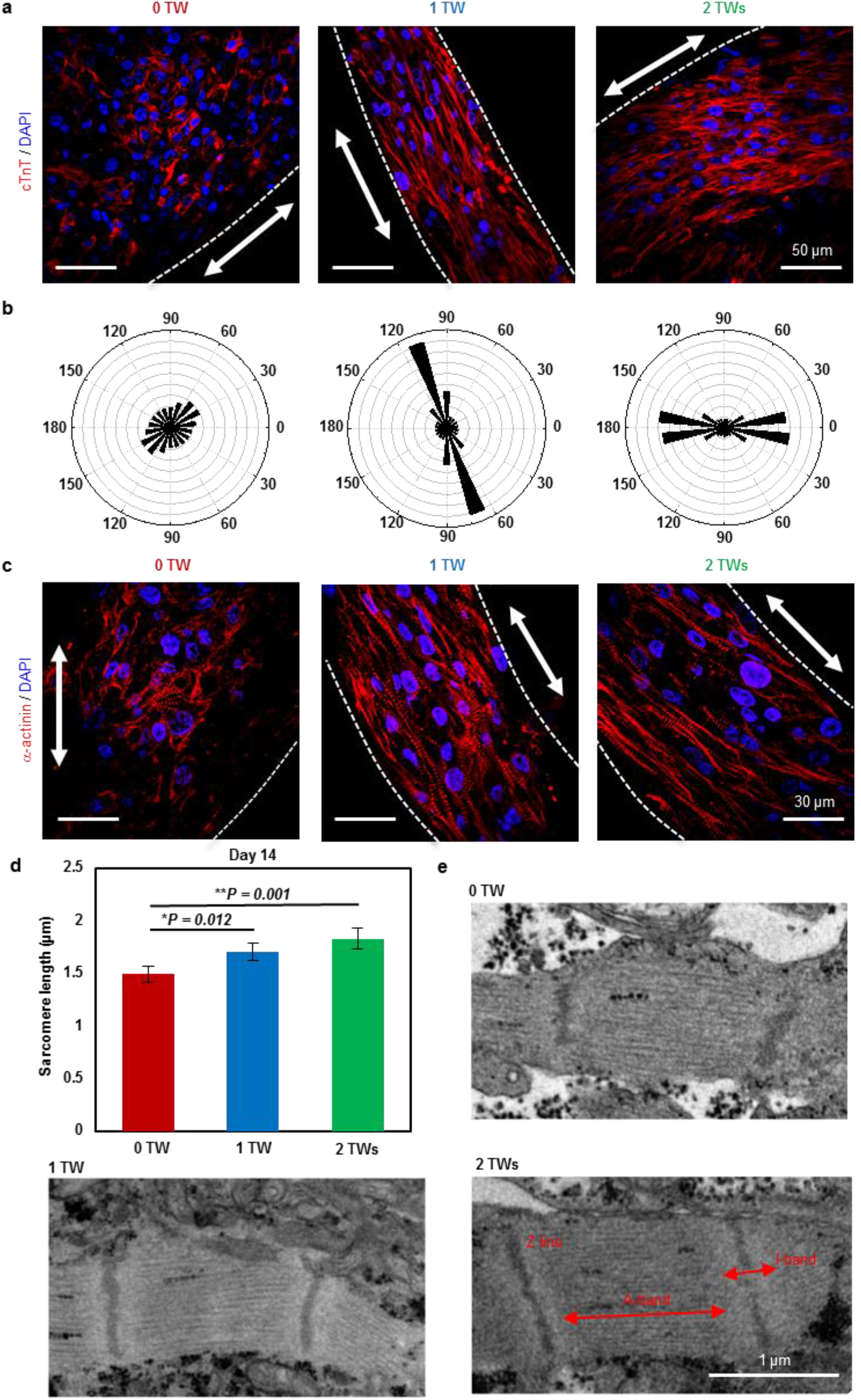
TWs promote CM alignment. (a) Representative confocal images of SOTRs with zero, one, or two TWs. CMs were stained with anti-TnT2 (red) and DAPI (blue), revealing CM alignment in TW samples. Arrows mark the ring orientation. (b) Angular graph based on Fourier component analysis of TnT2 orientation distribution. (c) Representative confocal images of stained SOTRs with different TWs (red: a-actinin; blue: DAPI). Arrows mark the ring orientation. (d) Sarcomere lengths of CMs within SOTRs on day 14 [mean ± s.d.; 0 TWs (n = 5); 1 TW (n = 5); 2 TWs (n = 4)]. \**P* < 0.05; \*\**P* < 0.01. (e) TEM analysis of SOTRs on day 14.

### TWs improve bioenergetics and Ca^2+^-handling properties

An extracellular-flux analyzer was used to characterize mitochondrial function in CMs within SOTRs. To record the maximum activity of the electron-transport chain from adenosine triphosphate (ATP) synthesis, mitochondrial ATP synthase was inhibited by oligomycin, and a proton-gradient discharger was added to measure maximum mitochondrial respiration. Our results indicated that the maximum respiration rate of groups with two TWs was significantly higher than that observed in groups with zero or one TW (Fig. 5a and b), indicating that a higher number of TWs led to increased mitochondrial activity.

**Figure 5.**
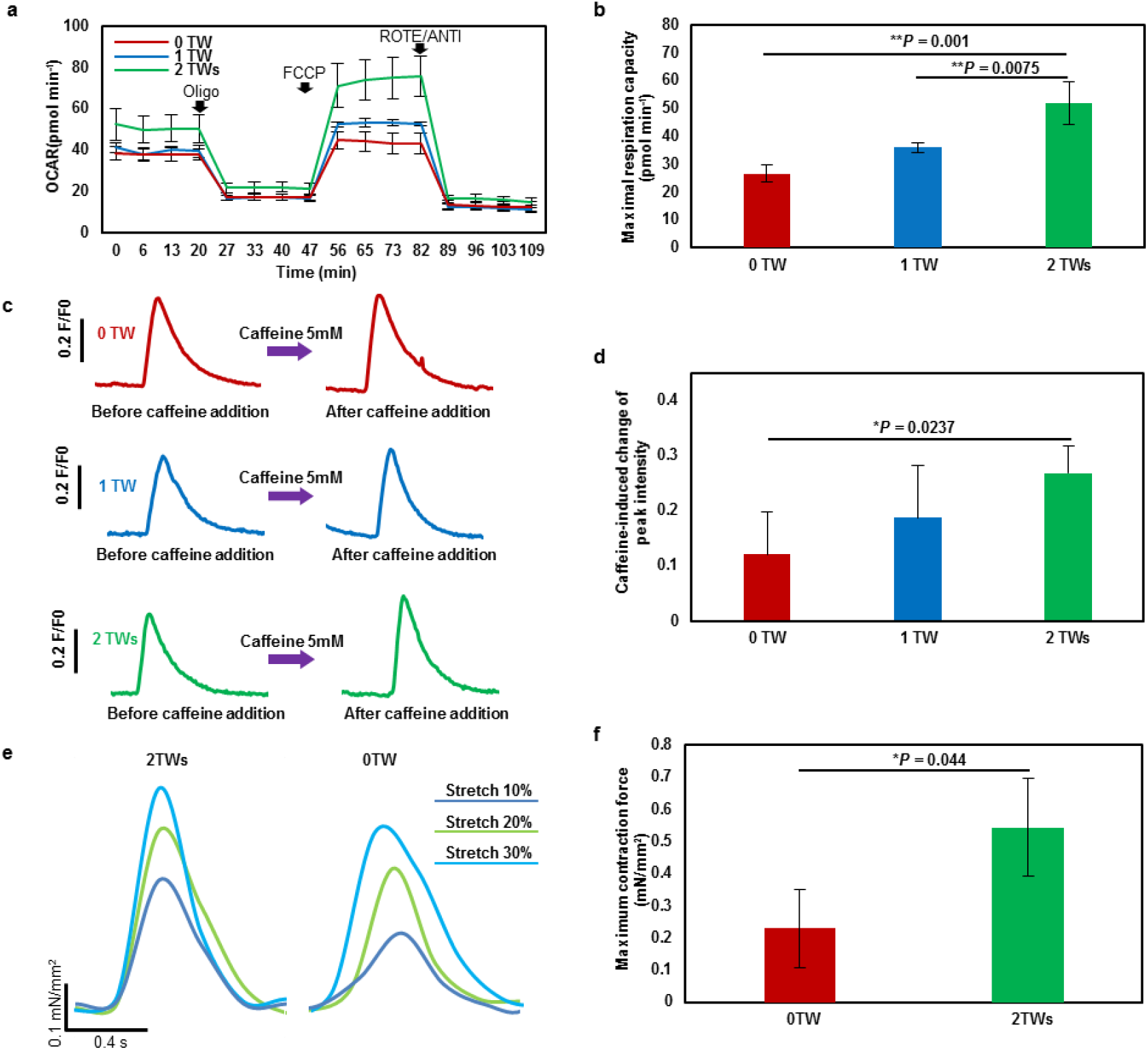
TWs improve Ca^2+^-handling properties. (a) Results of OCR assays. Olig, oligomycin; FCCP, carbonyl cyanide-4-(trifluoromethoxy) phenylhydrazone; ROTE/ANTI, rotenone and antimycin A. (b) Quantification of maximal respiration capacity (mean ± s.d.; *n* = 4). \*\**P* < 0.01. (c) Representative traces of Ca^2+^ transient following administration of 5 mM caffeine to CMs in SOTRs with zero TWs, one TW, and two TWs at day 7. SOTRs were immersed in room temperature medium to stop TWs on day 6. (d) Caffeine-induced change in peak intensity in different groups (mean ± s.d.; *n* = 6). \**P* < 0.05. (e) Force traces of SOTRs during progressive stretching. (f) Maximum contractile force of SOTRs (mean ± s.d.; *n* = 3). *\*P* < 0.05.

Because human CMs rely on Ca^2+^ influx from the sarcoplasmic reticulum (SR) for contraction, we investigated SR function by applying caffeine to SOTRs in order to open SR ryanodine channels^3^. Although all groups responded to the addition of 5 mM caffeine (Fig. 5c and d), SOTRs with two TWs exhibited significantly higher-intensity changes than groups with zero TWs, indicating a higher maturation level in Ca^2+^-handling properties and agreeing well with gene-expression results. Additionally, we recorded the contractile force of SOTRs using a mechanical tester (Fig. 5e and f), revealing that SOTRs with either zero or two TWs generated active forces that increased along with an applied stretching force in a Frank–Starling-like fashion^40^. Moreover, the group with two TWs generated forces with larger amplitude than that in the group with zero TWs (2 TWs: 0.54 ± 0.15 mN/mm^2^; 0 TW: 0.23 ± 0. 12 mN/mm^2^).

## Discussion

Excitation-contraction coupling is important for CM development and function^41^, and beating rate might play an important role in cardiac maturation. The human fetal heartbeat increases linearly during embryonic development from ~1 Hz to ~3 Hz within 9 weeks *in utero*^42,43^. hiPSC-CMs functionally and morphologically resemble fetal CMs; however, the average beating rate of hiPSC-CMs is much lower than human fetal CMs. Previous studies promoted CM maturation by increasing their beating rates; however, high beat rates of hiPSC-CMs could only be achieved by applying high frequency electrical stimulation^3,13,15^, sustainable application of which is technically challenging and may cause potential side effects, including pH shift, ROS generation, electrolysis^19^, and cell damage^17^.

In the present study, we created SOTRs with spontaneously generated TWs capable of sustainably causing CMs to beat at high frequencies (up to 4 Hz) in the absence of external stimulation. We found that CMs trained with TWs demonstrated dramatically improved structural maturation, enhanced cardiac gene expression, and Ca^2+^-handling properties in a frequency dependent manner. Because this activity might also cause hypoxia in SOTRs, in the future, an enhanced oxygen supply, such as that afforded by dynamic culture^5^, can be utilized during SOTR culture. Additionally, although the CMs trained by TWs remained less mature relative to those generated by state-of-art methods^10,13,44^, the maturation level afforded by TWs could be further improved by changing the stimulation window/duration and improving the beating frequency.

SOTRs were formed through a one-step, gel-free method utilizing a polydimethylsiloxane (PDMS) mold and commercially available Petri dishes. SOTR formation required <2 days, much shorter than the previously reported tissue-ring formation time (~7–14 days)^21,45^ due to multiple factors, including the use of gel-free medium, the low-attachment surface of the culture dish, and the ability of the CMs to aggregate into a 3D construct. This new methodology could be an important complement to current widely used methods^3,19,21,25^, and the rapid formation of the ring organoid might be important for future mass production for drug-screening applications.

Given the large cell number required for SOTR formation in one well (4 × 10^5^ cells), it remains difficult to achieve high-throughput assessment similar to those previously reported creating 200 tissue constructs per million cardiac cells^46^. Further optimizations are needed to scale down the PDMS mold and reduce the plated cell number. Additionally, because force generation is the primary function of CMs, force quantifications have been recently utilized to assess drug-related effects^46,47^; therefore, future mold design for SOTR formation might include a force transducer^48^, such as an elastic silicone pillar, capable of offering multiple parameters for more relevant assessment of drug response to pharmacologic agents. Furthermore, since the electrical pacing could be controlled with different loading level (by gradually increasing or decreasing frequency, or by tuning on/off region of stimulation over time)^13,46^, it would be useful to develop SOTR with controllable stimulation. SOTRs with controllable silicone pillar could be used to regulate the frequency of TWs in real-time.

It remains an open question as to what degree of CM maturation is necessary for regenerative applications^49^, such as injection into or engraftment onto an infarcted heart. The various maturation levels of SOTRs under different TW-training levels might offer preconditioned CMs for delivery into an infarcted myocardium, thereby allowing studies of how CM maturation affects therapeutic efficacy.

The key factors necessary for the sustained TWs were the closed-loop topology and the electric excitability of the enclosed cells. It could be that SOTRs with TWs can be created using other electrically active cells, such as neurons and retina cells. The looped TW might be useful for promoting the maturation of these cell types, which have previously been matured by electrical stimulation^50–53^. Moreover, because the frequency of the TWs within the SOTRs can be adjusted by changing SOTR diameter, and given that the TWs could be sustained for >89 days, it might be possible to create pacemaker tissue with an adjustable beat rate for use as an *in vitro* model for drug assessment or potentially for *in vivo* heart pacing.

In conclusion, we found that TWs could be spontaneously generated and maintained within a SOTR comprising hiPSC-CMs, and that the TWs were able to make the CMs beat at a high frequency comparable to that found *in utero* during embryonic development. Moreover, the TWs promoted frequency dependent structural and functional maturation of the CMs, thereby offering a supplementary approach to electrical-stimulation-based maturation of electrically active cell types.

## Methods

### Differentiation and culture of hiPSC-CMs

GCaMP3-positive hiPSCs (253G1) were cultured^54^ and differentiated^55^ according to previously published methods^54,56^. All experiments involving the use of hiPSCs were performed following Kyoto University and Osaka University guidelines. After 30 to 50 days of differentiation, CM colonies floating in medium were collected and dissociated into a single-cell suspension by stirring for between 1 h and 2 h in protease solution [0.1% collagenase type I, 0.25% trypsin, 1 U/mL DNase I, 116 mM NaCl, 20 mM HEPES, 12.5 mM NaH_2_PO_4_, 5.6 mM glucose, 5.4 mM KCl, and 0.8 mM MgSO_4_ (pH 7.35)]^56^. The CM purity was characterized with Flow Cytometry and the CMs with high purity (> 85 %) were used for following experiments.

### Device manufacture

PDMS (SYLGARD 184; Dow Corning, Midland, MI, USA) blocks were punched using a tissue puncher (Thermo Fisher Scientific, Waltham, MA, USA) to an inner diameter of 8 mm. After removal of the redundant central pillar, the remaining PDMS wells were collected, and PDMS wells and pillars with different diameters were centrally aligned and attached to the bottom of 24-well ultra-low attachment plates (Corning, Corning, NY, USA) as shown in Supplementary Fig. 1c. For micro-electrode array (MEA; Multi Channel Systems, Reutlingen, Germany) recording, the PDMS well and pillars were mounted to the MEA surface using silicone glue (Shin-Etsu Chemical, Tokyo, Japan) to fix the well and pillar to the bottom of the well. After drying and treatment with UV light for 30 min, the PDMS wells were ready for CM culture.

### SOTR generation

Before cell seeding, single CMs were filtered using a 40-μm cell strainer (BD Falcon; Becton Dickenson, Franklin Lakes, NJ, USA) and resuspended at a density of 2 × 10^6^ cells/mL in serum-supplemented cardiac differentiation medium (Sigma-Aldrich, St. Louis, MO, USA) containing 20% fetal bovine serum (FBS; Gibco, Grand Island, NY, USA), 1% minimum essential medium non-essential amino acid solution (Sigma-Aldrich), 1% penicillin-streptomycin (Gibco), 2 mM L-glutamine (Sigma-Aldrich), 0.001% 2-mercaptoethanol (Gibco), and 0.005 N NaOH along with 10 ng/mL bone morphogenetic protein 4 (R&D Systems, Minneapolis, MN, USA). 4 × 10^5^ cells were plated in each PDMS culture well with 3-mm pillar. For wells with other diameters, the cell number was fixed to 4 × 10^5^ or proportional to the diameter. After plating, CMs settled in the wells, aggregated, and congregated around the central pillar to form densely packed tissue rings within 2 days. The medium was changed to serum-free medium (cardiac differentiation medium without FBS) from day 2. After that, fresh medium was changed every 4 days. Before medium changing, the fresh medium was pre-warmed to 37 degree. The dishes were placed on top of a prewarmed metal block during medium changing. The medium was pipetted gently and slowly into the wells.

### Optical mapping and measurement

Progression of tissue assembly was observed using a fluorescence microscope (Olympus IX71; Olympus, Tokyo, Japan) equipped with a charge-coupled device (CCD) camera (Exiblue; Qimaging, Surrey, BC, Canada; or DP74; Olympus). GCaMP3 was excited from 450 nm to 490 nm, and fluorescence images of GCaMP3-positive CMs were recorded with 8 × 8 binning of CCD pixels at 30 frames/s. Images were processed to obtain data using ImageJ (NIH, Bethesda, MD, USA) and MATLAB (R2014b; MathWorks, Natick, MA, USA) using a customized program. Activation-time mapping was performed using a custom plug-in in ImageJ, as previously described^57^.

### Transient and electrophysiological Ca^2+^ characterization

Ca^2+^ transience was recorded by fluorescence imaging of GCaMP3-positive CMs with the same experimental setup as that described for optical mapping and measurement. To characterize the spontaneous beating of SOTRs, TWs were removed from the SOTRs 1 day before recording by emerging the SOTRs into medium at room temperature for 1 min. The L-type Ca^2+^-channel blocker verapamil (1 mM; Sigma-Aldrich) and 5 mM caffeine (Wako Pure Chemical Industries, Ltd. Osaka, Japan) were added directly to the chamber containing the SOTRs during imaging, as described in the figure legends.

Extracellular recording of field potentials (FPs) was performed using an MEA data-acquisition system (USB-ME64; Multi Channel Systems). Signals were recorded from days 5 and 6 after CM seeding, and data were collected and processed using MC_Rack (Multi Channel Systems) or LabChart (ADInstruments, Dunedin, New Zealand). The amplitude, QT interval, and beat rate were determined by analyzing the wave form of the FP.

### Histology

SOTRs were rinsed three times with phosphate-buffered saline (PBS), fixed in 4% paraformaldehyde (PFA) in PBS for 30 min, and embedded in paraffin. Thin sections were sliced and stained with HE (Muto Chemical Corporation, Tokyo, Japan), and observation was carried out using a CKX41 microscope (Olympus).

### Immunostaining and imaging

SOTRs were fixed in 4% PFA at room temperature for 30 min, permeabilized with 0.5% (v/v) Triton X-100 in Dulbecco’s (D)-PBS at room temperature for 1 h, and incubated in blocking solution [0.1% (v/v) Tween-20, 5% (v/v) normal donkey serum, 3% (v/v) bovine serum albumin, and 5% (v/v) normal goat serum in D-PBS] at 4°C for 16 h. SOTRs were then incubated with the primary antibodies anti-troponin T2 (TnT2; mouse monoclonal IgG; 1:200; SC-20025; Santa Cruz Biotechnology, Dallas, TX, USA), anti-α-actinin (mouse monoclonal IgG, 1:1000; A7811; Sigma-Aldrich), or anti-β-MHC (mouse MYH7 monoclonal IgM, 1:100; SC-53089; Santa Cruz Biotechnology) at 4°C overnight. SOTRs were then rinsed with PBS and incubated with secondary antibodies diluted 1:300 in blocking buffer (Alexa Fluor 594 anti-mouse IgG; 715-586-150; and DyLight-594 anti-mouse IgM; 715-516-020; Jackson ImmnoResearch, West Grove, PA, USA) at room temperature for 1 h. 4’-6-Diamidino-2-phenylindole (DAPI; 300 nM; Wako Pure Chemical Industries, Ltd.) was used to counterstain nuclei at room temperature for 30 min, after which images were captured using a confocal microscope (FV1200; Olympus). CM orientation within SOTRs was determined using the Fourier component analysis plugin “Directionality” in ImageJ^47^ (NIH) that calculated the orientation distribution for the red color channel. Fluorescence quantification of β-MHC was performed by calculating the gray value averaged over the area in all sample groups, with all the fluorescence values were normalized to that of zero TW group.

### Transmission electron microscopy (TEM)

SOTRs were washed with 0.1 M Sorenson’s buffer (pH 7.4) twice and fixed with 2.5% glutaraldehyde (Sigma-Aldrich) in 0.1 M Sorenson’s buffer (pH 7.4), after which samples were post-fixed with 1% OsO4 in Sorrenson’s buffer. The samples were then embedded, sliced, and stained with lead citrate and examined under a JEOL1010 transmission electron microscope (JEOL Ltd., Tokyo, Japan).

### Mitochondrial respiration assay

Mitochondrial function was analyzed using a Seahorse XF96 extracellular flux analyzer (Agilent Technologies, Carlsbad, CA, USA). After a 14-day culture, SOTRs were dissociated into a single-cell suspension by stirring for 30 min in protease solution, followed by cell seeding onto a cell-culture microplate (Agilent Technologies) at a density of 20,000 cells per XF96 well. OCR assays were performed 3 days after seeding, and culture medium was changed to the base medium (Seahorse XF assay media supplemented with 1 mM sodium pyruvate; Life Technologies, Carlsbad, CA, USA). Substrates and inhibitors were injected during measurements at a final concentrations of 3.5 μM 4-(trifluoromethoxy) phenylhydrazone (FCCP; Seahorse Bioscience, Billerica, MA, USA), 1 μM oligomycin, 0.5 μM antimycin, and 0.5 μM rotenone for the MitoStress assay.

### Flow Cytometry

Before used for generating SOTRs, hiPSCs-CMs fixed in 4% PFA at room temperature for 30 min, permeabilized with 0.5% v/v Triton X-100 in Dulbecco’s (D)-PBS at room temperature for 30 min, incubated with anti-cTnT antibodies (mouse monoclonal IgG, 1:200; Santa Cruz Biotechnology: SC-20025) or isotype-matched antibodies (BD Phosphoflow: 557782) at 37 °C for 30 min, washed with D-PBS, and then incubated with Alexa Fluor 488 anti-mouse IgG (1:500; Jackson ImmnoResearch: 715-546-150). Cells were then washed twice with D-PBS and analyzed using a FACS Canto II flow cytometer (BD Biosciences, USA) and the FlowJo software (Treestar Inc., USA). Data shown are representative of five independent experiments.

### Contractility analysis

SOTR contractility was measured using a micron-scale mechanical-testing system (MicroSquisher; CellScale Biomaterials Testing, Waterloo, ON, Canada). TWs were stopped by exchanging medium at room temperature 1 day before measurement, and the SOTR was removed from the pillar and fixed on a stage. A cantilever beam with a diameter of 0.30 mm was pressed onto the SOTR, with the force calculated by cantilever-beam deflection in response to differential displacement. All samples were tested at 37°C.

### qPCR

Total RNA in SOTRs was harvested using Trizol reagent (Life Technologies) according to manufacturer instructions, and RNA concentration was determined using a NanoDrop1000 spectrophotometer (Thermo Fisher Scientific). cDNA was synthesized with a first-strand synthesis kit (TaKaRa, Shiga, Japan) and analyzed by qPCR using SYBR Green PCR master mix (Life Technologies) and the qBiomarker validation PCR array (IPHS-102A; Qiagen, Hilden, Germany) in a 96-well format according to manufacturer instructions. Cycling conditions were set as follows: initial denaturation at 95°C for 10 min, followed by 40 cycles at 95°C for 15 s and 60°C for 70s. Reactions were performed in a StepOnePlus real-time PCR system (Life Technologies). Gene expression was determined using the 2^-ΔΔCt^ method and relative to *glyceraldehyde 3-phosphate dehydrogenase* expression. Heatmaps were created by the “pheatmap” package (https://cran.r-project.org/web/packages/pheatmap/index.html), and clustering order was produced using the Ward.D clustering algorithm (https://stat.ethz.ch/R-manual/R-devel/library/stats/html/hclust.html).

### RNA quantification and qualification for RNA sequencing

RNA degradation and contamination was monitored on 1% agarose gels. RNA purity was determined using a NanoPhotometer spectrophotometer (Implen, Westlake Village, CA, USA). RNA concentration was measured using a Qubit RNA assay kit in a Qubit 2.0 fluorometer (Life Technologies). RNA integrity was assessed using the RNA Nano 6000 assay kit and the Agilent Bioanalyzer 2100 system (Agilent Technologies).

### Library preparation for RNA sequencing

A total of 1.5 μg RNA per sample was used as input material for RNA sample preparations. Sequencing libraries were generated using a NEBNext UltraTM RNA library prep kit Illumina (New England Biolabs, Ipswich, MA, USA) according to manufacturer instructions, and index codes were added to attribute sequences to each sample. Briefly, mRNA was purified from total RNA using poly-T oligo-attached magnetic beads, and fragmentation was performed using divalent cations under an elevated temperature in NEBNext First Strand Synthesis reaction buffer (5x; New England Biolabs). First-strand cDNA was synthesized using a random hexamer primer and M-MuLV reverse transcriptase (RNase H-; New England Biolabs). Second-strand cDNA synthesis was subsequently performed using DNA polymerase I and RNase H. Remaining overhangs were converted into blunt ends via exonuclease/polymerase activities. After adenylation of the 3’ ends of the DNA fragments, NEBNext Adaptor (New England Biolabs) with a hairpin loop structure was ligated to prepare for hybridization. To select cDNA fragments of the correct length, the library fragments were purified using the AMPure XP system (Beckman Coulter, Beverly, MA, USA), after which 3 μL USER enzyme (New England Biolabs) was incubated with size-selected, adaptor-ligated cDNA at 37°C for 15 min, followed by 5 min at 95°C. PCR was performed using a Phusion high-fidelity DNA polymerase, universal PCR primers, and an Index (X) primer (New England Biolabs). Products were purified (AMPure XP; Beckman Coulter), and library quality was assessed using the Agilent Bioanalyzer 2100 system (Agilent Technologies).

### Clustering and sequencing

Clustering of the Index-coded samples was performed on a cBot cluster generation system using a HiSeq 4000 PE cluster kit (Illumia, San Diego, CA, USA) according to manufacturer instructions. After cluster generation, the library preparations were sequenced on an Illumina Hiseq 4000 platform (Illumina), and 150-bp paired-end reads were generated.

After quality control, the paired-end reads were aligned to the hg19 human reference genome (UCSC; https://genome.ucsc.edu/) using TopHat (v2.0.12; https://ccb.jhu.edu/software/tophat/index.shtml)^48^. We quantified the relative expression level as log2(FPKM + 1) to perform PCA analysis and hierarchical clustering. Uniquely mapped reads counted by HTSeq^49^ were used to conduct differential-expression analysis in DEseq2^50^. A total of 349 gens (p<0.05, fold change>1.5) were selected for the GO analysis. We plotted heatmaps using the “pheatmap” package in R (https://cran.r-project.org/web/packages/pheatmap/index.html) and performed GO enrichment analysis using Metascape^58^ (http://metascape.org/).

### Statistical analysis

All quantitative data are presented as the mean ± standard deviation of the mean (mean ± s.d.). The differences between experimental groups were analyzed by Student’s *t* test (between two groups) or one-way analysis of variance, followed by Tukey’s post hoc test (among three groups). A P < 0.05 was considered statistically significant.

## Supplementary Information

### Circulating traveling waves enhance the maturation of hiPSC-derived cardiomyocytes in self-organized tissue rings

#### Mathematical model

We constructed a mathematical model to simulate the phenomena observed in experiments. This model comprises two critical parts: 1) modeling of the action potential (AP) in a cell and coupling between two neighboring cells. First, when we simulated the AP of a single human induced pluripotent stem cell-derived cardiomyocyte (hiPSC-CM), we referenced the classic Priebe & Beuckelmann (PB) model^1^ of human ventricular AP based on the Hodgkin-Huxley formalism, with the equations derived from experimental data. To increase computational efficiency and stability, we referenced Panfilov’s reformulated model^2^ based on the PB model. Unlike human ventricular cells, hiPSC-CMs exhibit an ability to beat spontaneously, indicating that hiPSC-CMs are similar to sinoatrial node cells. Therefore, we added hyperpolarization-activated current^3^, I_f_, to reformulate the PB model. There was evidence that hiPSC-CMs differed significantly from adult ventricular CMs according to their reduced inward rectifier K^+^ currents (I_k1_) and the presence of prominent pacemaker currents (If) that contribute to hiPSC-CM automaticity^4^. Therefore, we changed the maximum conductance value, g_k1_, of the inward rectifier K^+^ currents (I_k1_) in our mathematical model. A detailed description of the equations governing were described previously^1–3^. Second, there exists a conduction of electrical signals between neighboring cells. Previous studies treated this process using cable equations^5^:

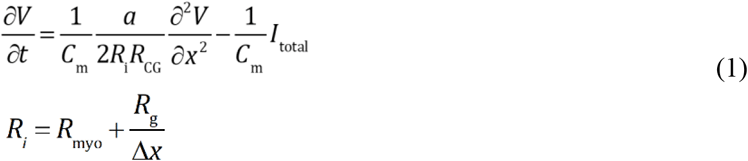

where *V* is the membrane potential, *C*_m_ is the membrane capacity, *I*_total_ is the sum of individual membrane ionic currents in a single cell, *a* is the cell radius, Δ*x* is the length of a single cell, *R*_CG_ is the ratio between capacitive and geometric areas, and *R*_i_ comprises the myoplasmic resistance, *R*_myo_, and gap-junction resistance, *R*_g_, as shown. By combining and modifying the above parts, we established a mathematical model of coupled hiPSC-CMs. To create the model in agreement with our experimental results, we set parameters as follows: g_Na_ = 8 nS/pF, g_to_ = 0.35 nS/pF, g_K_ = 0.1 nS/pF, g_K1_ = 1.17 nS/pF, g_f_ = 0.08 nS/pF, *R*_g_ = 0.1Ω, and *R*_myo_ = 0.007 Ω/um. We used N cells arranged next to each other in one dimension with periodic boundary conditions to simulate the self-organized tissue ring (SOTR) in the experiment.

#### The dependence of traveling wave (TW) speed on ring diameter

Both in the experiment and the simulation, TW speed increased along with ring diameter (Fig. 3d). A key determinant of TW speed was the difference in membrane potential between the yet-to-be-activated cell and the neighboring cell that activates it. All hiPSC-CMs are capable of spontaneously beating, and their APs change periodically. Therefore, maintaining spontaneous beating ability requires the Funny current (hyperpolarization-activated current^3^), If, which helps resting cells gradually elevate their APs beyond a threshold, followed by the initiation of depolarization. When stable TWs appear in a ring, the time between two successive beats becomes shorter than that of its own spontaneous beat rate. This is because in the case of TWs, the beating of a cell is due to activation by its neighbor, which would occur prior to its self-activation by the Funny current. The larger the ring diameter, the longer it will take the TW to traverse it, and the longer a cell waits to be activated by its neighbor.

This will bring the cell closer to its own self-activation point, resulting in a more rapid activation when the neighbor is activated and a more rapidly propagating TW. The above intuitive argument can be quantitatively expressed in the following equations.

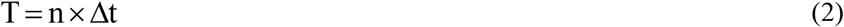

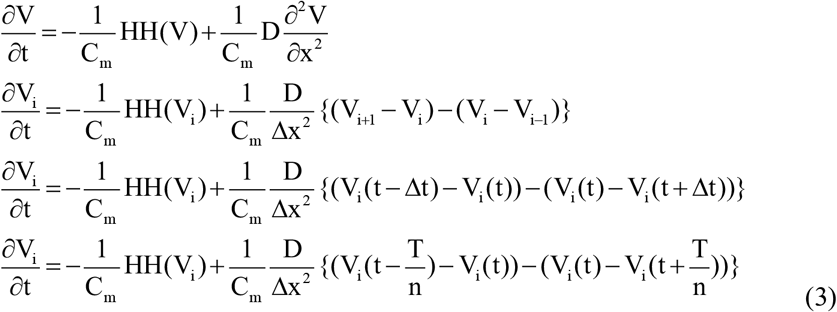

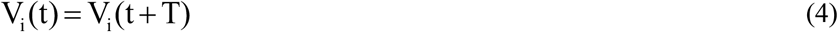

T: the time for one TW to travel around the ring

n: the number of cells

Δt: the difference in time between the point at which the electrical signal arrives at two adjacent cells

V: membrane potential

V_i_: the membrane potential of the i^th^ cell

HH(V): derivation of V and I using the HH equation

D: diffusion coefficient

Given the diameter of the ring, the number of cells is determined (n). Using the periodic boundary condition (Eq. 4), the above equations can be solved to find a relationship between the cycle time of the membrane potential of a single cell and the diameter of the ring (Supplementary Fig. 4a).

To verify the accuracy of the above explanation, we performed two simulations. First, the Funny current resulted in different AP differences between the two adjacent cells at the wave front in rings with different diameters. Therefore, we deleted the Funny current in every cell, resulting in the loss of spontaneous beating ability and an inability of its AP to change in the resting state over time. This meant that whenever a cell was activated, the difference in AP between the activating cell and the activated cell would be the same, regardless of the diameter of the ring. In this simulation, we found that the diameter of the ring did not affect TW velocity, suggesting that transmitted electrical signals in different ring diameters had no effect on speed due to their shared difference in electrical potential (Supplementary Fig. 4b).

Second, we artificially raised the minimum membrane potential of cells in the resting state and maintained their excitable characteristics (with no Funny current). Unlike the first simulation, we established gradients of differences in electrical potential between activating and activated cells. After measuring the propagation velocity of the TWs, we found that the propagation speed decreased as the difference in electrical potential increased (Supplementary Fig. 4c).

These two simulations demonstrated that different TW speeds in rings of different diameters were caused by differences in the electrical potential between activated and activating cells. The smaller the ring diameter, the greater the difference in electrical potential and the smaller the propagation speed.

---

**Supplementary Figure 1.**
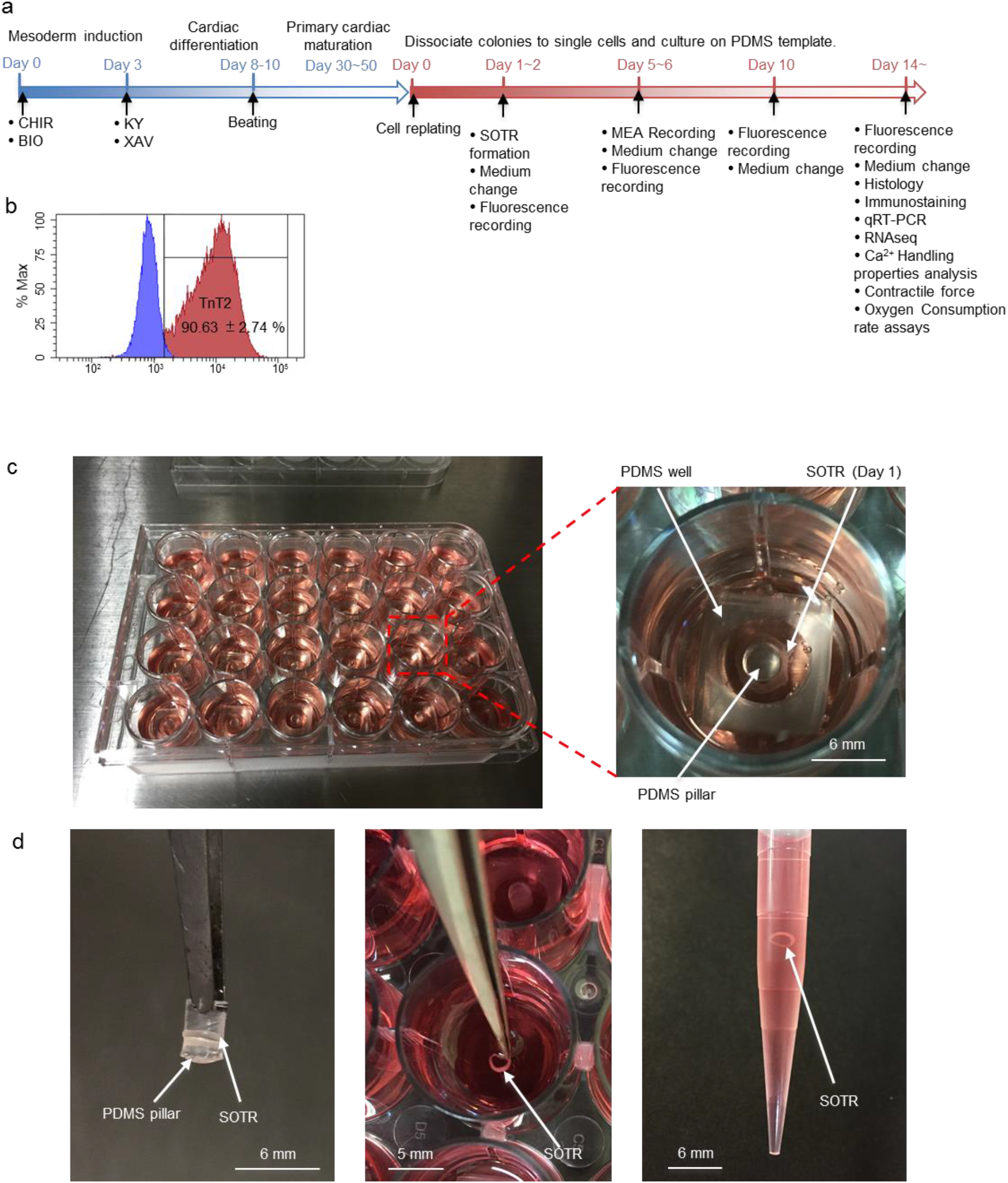
SOTR formation. (a) Schematic representation of CM differentiation and functional assessment in SOTRs. (b) Flow cytometric analysis of the CMs used for generating SOTRs. (mean ± s.d.; *n* = 4) (c) SOTRs in a 24-well culture dish. The enlarged figure shows the PDMS well and pillar, between which the cells were plated and organized into a tissue ring. (d) SOTR with a PDMS pillar on day 14. The SOTR could be removed from the pillar and transferred with a pipette.

**Supplementary Figure 2.**
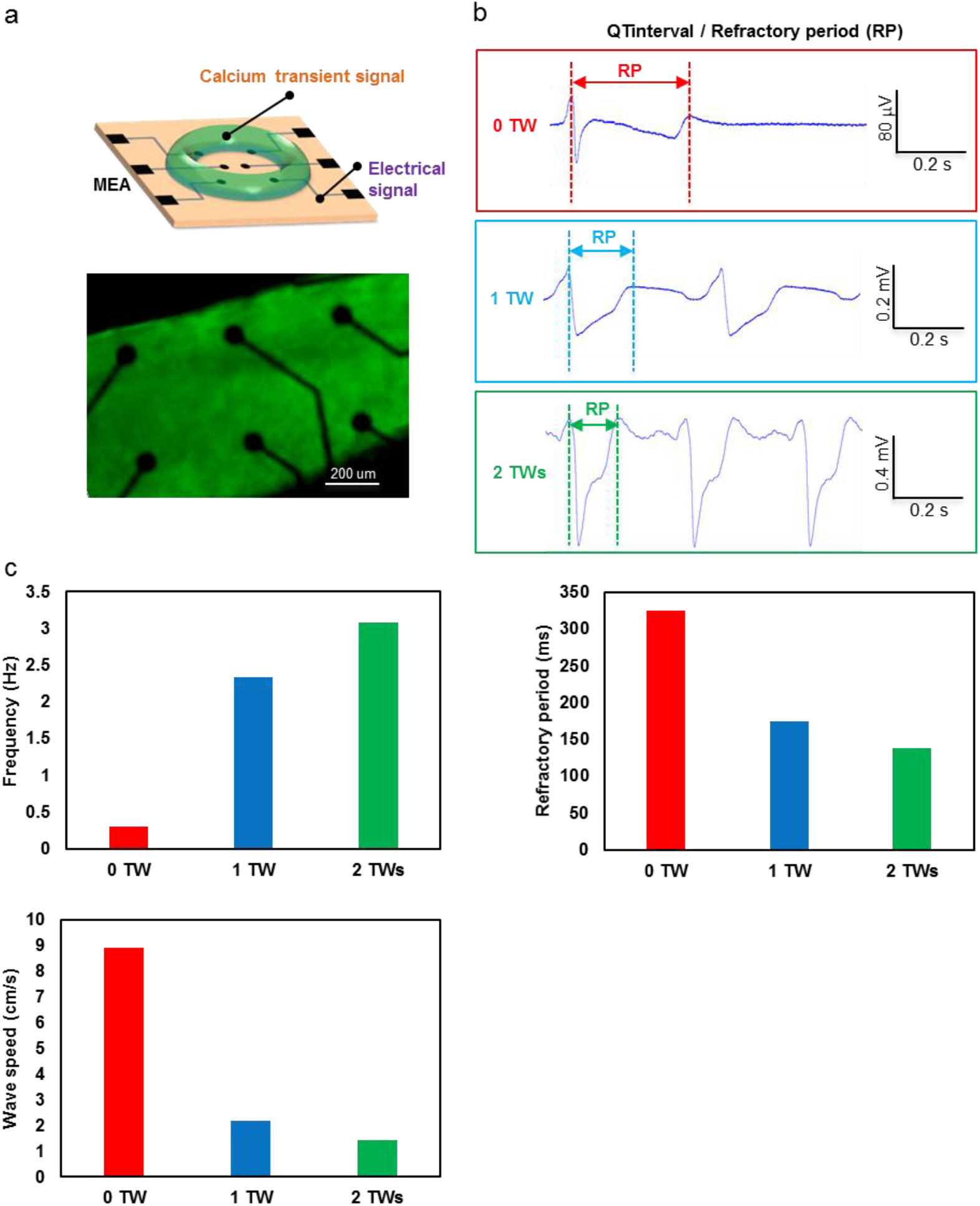
FP recording by micro-electrode arrays. a) Representative setup for FP recording by electrodes underneath a SOTR on day 6. See also supplementary video 5. (b) Representative FP recorded by an electrode underneath a SOTR with or without TWs. (c) Frequency, QT interval (correlated with AP duration and refractory period), and wave speed in SOTRs with or without TWs.

**Supplementary Figure 3.**
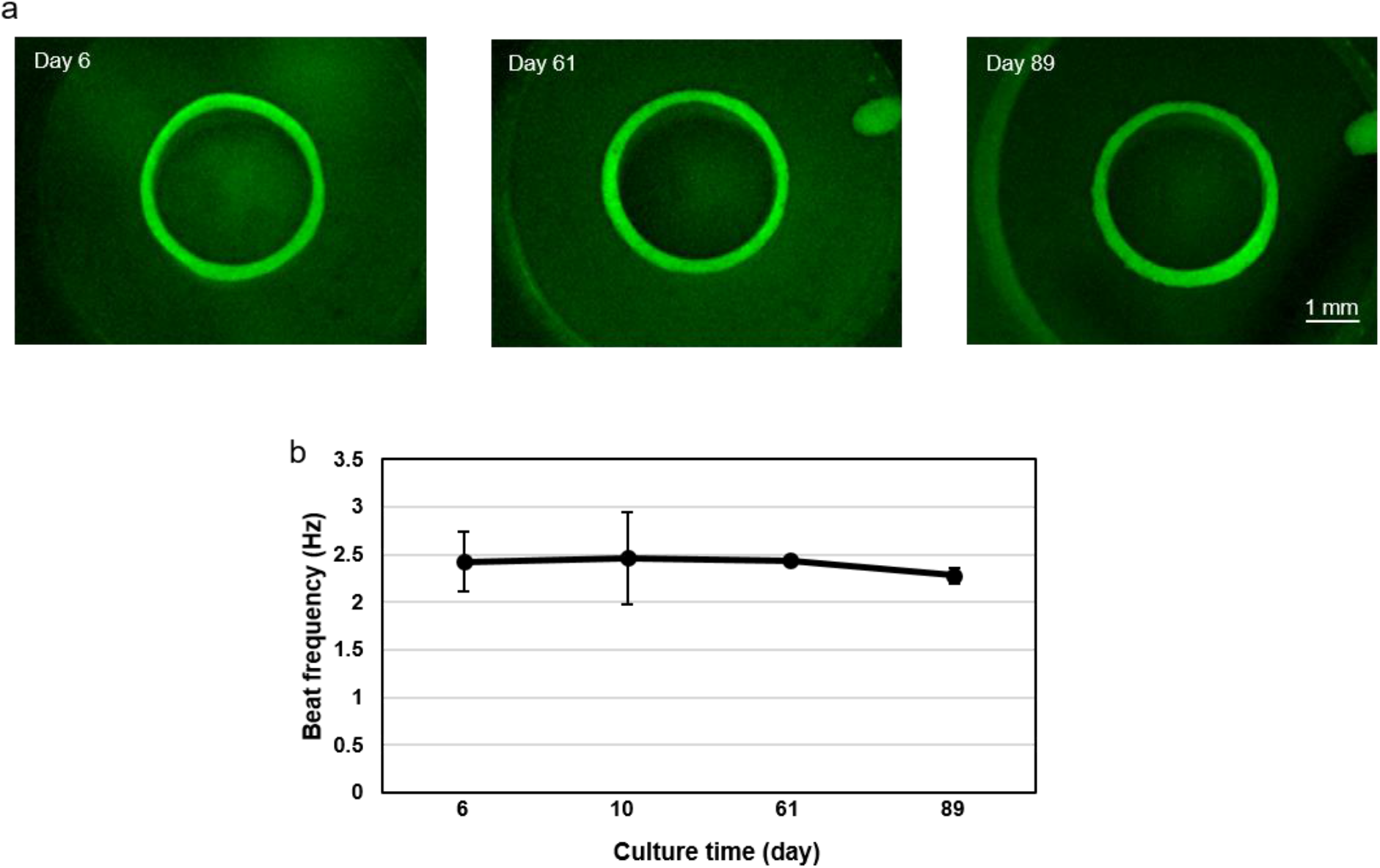
Maintenance of TWs in SOTRs for >89 days. (a) Fluorescence images of a SOTR with one TW at different culture times. (b) The beat rate (Hz) of SOTRs with one TW during a long-term culture (mean ± s.d.; *n* = 3).

**Supplementary Figure 4.**
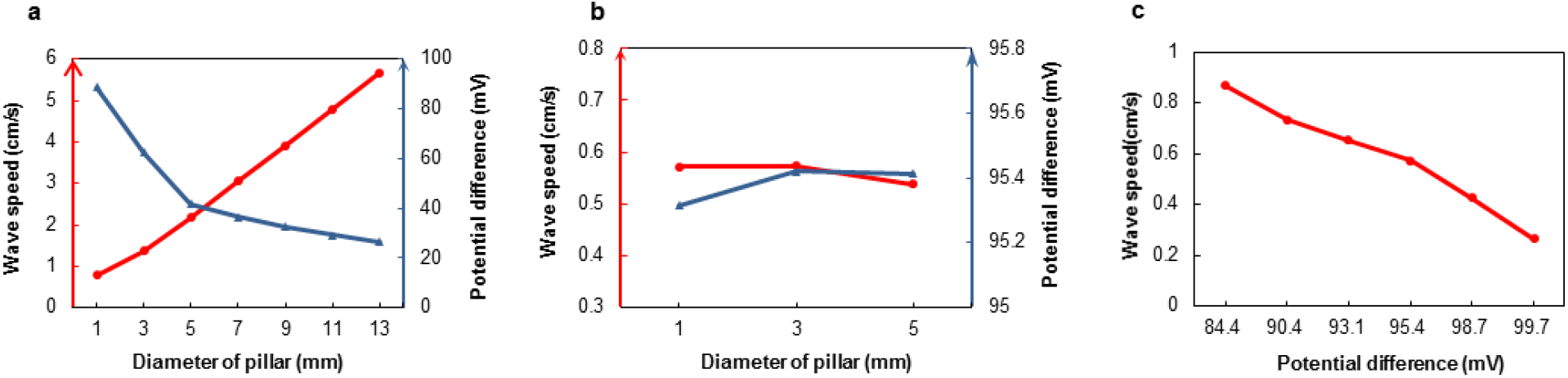
Relationships between TW speed and differences in membrane potential. (a) Relationships between TW speed, differences in electrical potential between neighboring cells just before activation (at the front tip of the wave), and ring diameter. The blue line represents the TW speed, and the orange line represents the difference in electrical potential. (b) The diameter of the ring does not affect TW velocity when cells lose the Funny current. The blue line represents TW velocity, and the orange line represents the difference in electrical potential. (c) Differences in electrical potential affect TW speed.

**Supplementary Figure 5.**
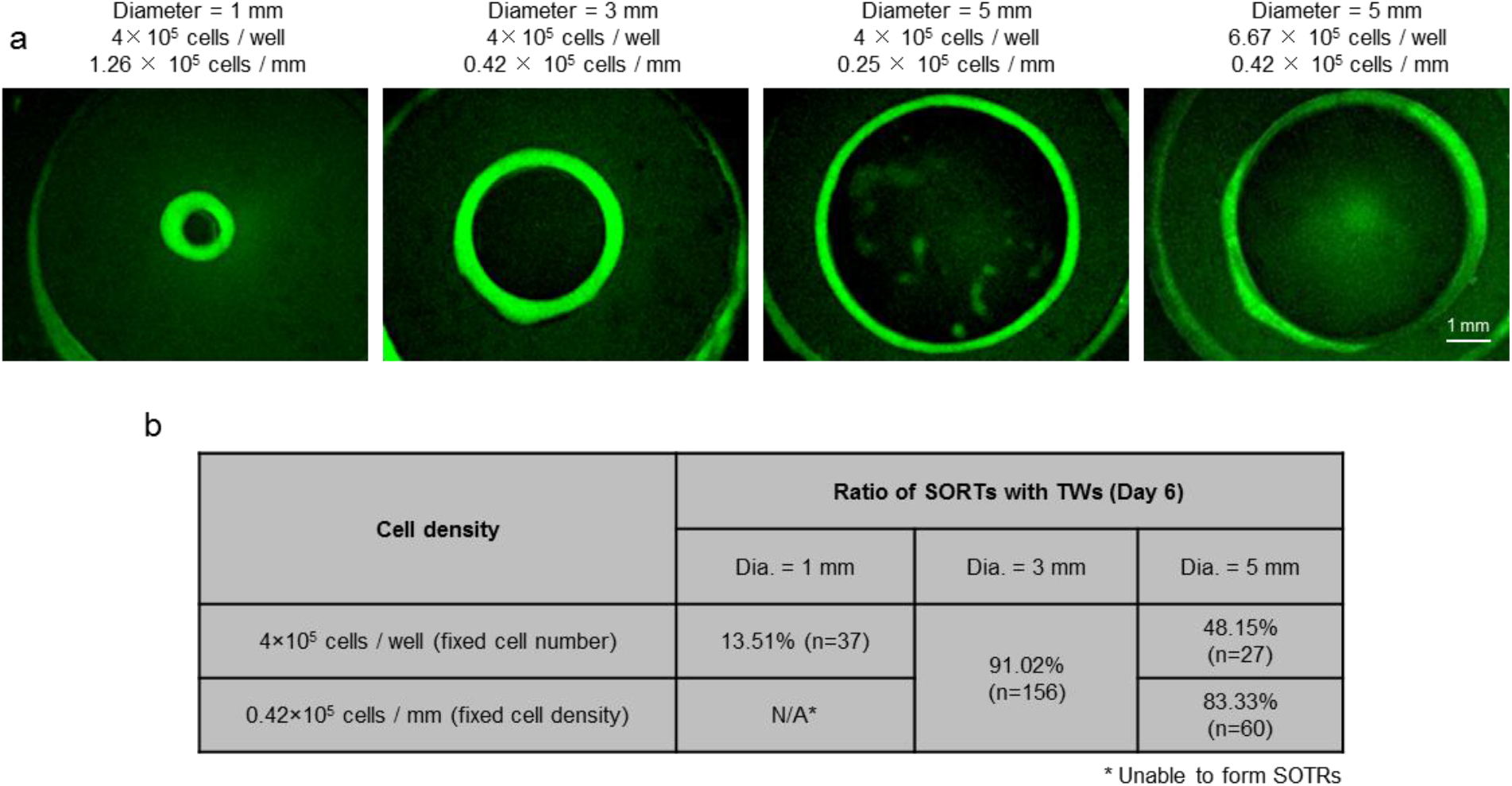
SOTR formation using pillars with different diameters. (a) Fluorescence images of GCaMP3-positive SOTRs with different diameters at day 6. (b) Percentage of SORTs with TWs on day 6 after cell plating with different pillar diameters.

**Supplementary Figure 6.**
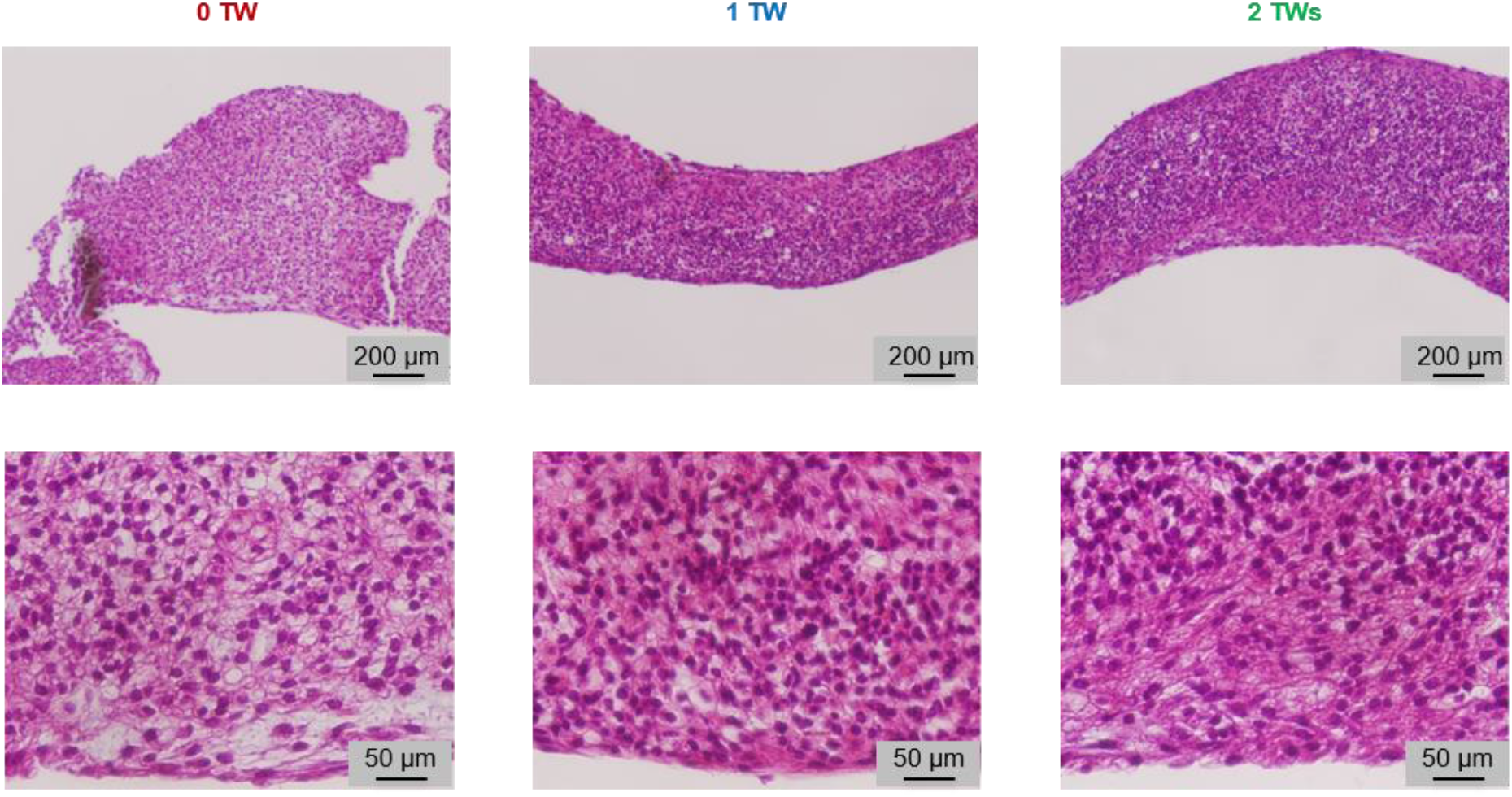
Histology of SOTRs with different TWs on day 6 after cell plating.

## Reference

1 Karakikes, I., Ameen, M., Termglinchan, V. & Wu, J. C. Human Induced Pluripotent Stem Cell–Derived Cardiomyocytes Insights Into Molecular, Cellular, and Functional Phenotypes. Circulation Research. 117, 80–88 (2015).

2 Dolnikov, K. et al. Functional properties of human embryonic stem cell–derived cardiomyocytes: intracellular Ca2+ handling and the role of sarcoplasmic reticulum in the contraction. Stem cells. 24, 236–245 (2006).

3 Nunes, S. S. et al. Biowire: a platform for maturation of human pluripotent stem cell–derived cardiomyocytes. Nat. Methods. 10, 781–787 (2013).

4 Davis, R. P. et al. Cardiomyocytes derived from pluripotent stem cells recapitulate electrophysiological characteristics of an overlap syndrome of cardiac sodium channel disease. Circulation. 125, 3079–3091 (2012).

5 Jackman, C. P., Carlson, A. L. & Bursac, N. Dynamic culture yields engineered myocardium with near-adult functional output. Biomaterials. 111, 66–79 (2016).

6 Ulmer, B. M. et al. Contractile Work Contributes to Maturation of Energy Metabolism in hiPSC-Derived Cardiomyocytes. Stem cell reports. 10, 834–847 (2018).

7 Shadrin, I. Y. et al. Cardiopatch platform enables maturation and scale-up of human pluripotent stem cell-derived engineered heart tissues. Nature communications. 8, 1825 (2017).

8 Gao, L. et al. Myocardial Tissue Engineering With Cells Derived From Human-Induced Pluripotent Stem Cells and a Native-Like, High-Resolution, 3-Dimensionally Printed Scaffold. Circulation Research. 120, 1318–1325 (2017).

9 Parikh Shan, S. et al. Thyroid and Glucocorticoid Hormones Promote Functional T-Tubule Development in Human-Induced Pluripotent Stem Cell-Derived Cardiomyocytes. Circulation Research. 121, 1323–1330 (2017).

10 Herron, T. J. et al. Extracellular matrix–mediated maturation of human pluripotent stem cell-derived cardiac monolayer structure and electrophysiological function. Circulation: Arrhythmia and Electrophysiology. 9, e003638 (2016).

11 Brevet, A., Pinto, E., Peacock, J. & Stockdale, F. E. Myosin synthesis increased by electrical stimulation of skeletal muscle cell cultures. Science. 193, 1152–1154 (1976).

12 McDonough, P. M. & Glembotski, C. Induction of atrial natriuretic factor and myosin light chain-2 gene expression in cultured ventricular myocytes by electrical stimulation of contraction. J. Biol. Chem. 267, 11665–11668 (1992).

13 Ronaldson-Bouchard, K. et al. Advanced maturation of human cardiac tissue grown from pluripotent stem cells. Nature. 556, 239–243 (2018).

14 Nunes, S. S. et al. Biowire: a platform for maturation of human pluripotent stem cell-derived cardiomyocytes. Nat Methods. 10, 781–787 (2013).

15 Eng, G. et al. Autonomous beating rate adaptation in human stem cell-derived cardiomyocytes. Nature communications. 7 (2016).

16 Tandon, N. et al. Electrical stimulation systems for cardiac tissue engineering. Nature protocols. 4, 155–173 (2009).

17 Merrill, D. R., Bikson, M. & Jefferys, J. G. R. Electrical stimulation of excitable tissue: design of efficacious and safe protocols. J. Neurosci. Methods. 141, 171–198 (2005).

18 Hirt, M. N. et al. Functional improvement and maturation of rat and human engineered heart tissue by chronic electrical stimulation. Journal of molecular and cellular cardiology. 74, 151–161 (2014).

19 Godier-Furnémont, A. F. et al. Physiologic force-frequency response in engineered heart muscle by electromechanical stimulation. Biomaterials. 60, 82–91 (2015).

20 van Meer, B. J., Tertoolen, L. G. & Mummery, C. L. Measuring physiological responses of human pluripotent stem cell derived cardiomyocytes to drugs and disease. STEM CELLS. 34, 2008–2015 (2016).

21 Zimmermann, W.-H. et al. Tissue engineering of a differentiated cardiac muscle construct. Circulation research. 90, 223–230 (2002).

22 Dhein, S. et al. Mechanical control of cell biology. Effects of cyclic mechanical stretch on cardiomyocyte cellular organization. Progress in biophysics and molecular biology. 115, 93–102 (2014).

23 Li, Y. et al. Chinese-Noodle-Inspired Muscle Myofiber Fabrication. Advanced Functional Materials. 25, 5999–6008 (2015).

24 Tulloch, N. L. et al. Growth of engineered human myocardium with mechanical loading and vascular coculture. Circulation research. 109, 47–59 (2011).

25 Mihic, A. et al. The effect of cyclic stretch on maturation and 3D tissue formation of human embryonic stem cell-derived cardiomyocytes. Biomaterials. 35, 2798–2808 (2014).

26 Shaheen, N. et al. Human Induced Pluripotent Stem Cell-Derived Cardiac Cell Sheets Expressing Genetically Encoded Voltage Indicator for Pharmacological and Arrhythmia Studies. Stem cell reports. 10, 1879–1894 (2018).

27 Kadota, S. et al. Development of a reentrant arrhythmia model in human pluripotent stem cell-derived cardiac cell sheets. European heart journal. 34, 1147–1156 (2012).

28 González, H., Nagai, Y., Bub, G., Glass, L. & Shrier, A. Reentrant waves in a ring of embryonic chick ventricular cells imaged with a Ca2+ sensitive dye. BioSyst. 71, 71–80 (2003).

29 Kawatou, M. et al. Modelling Torsade de Pointes arrhythmias in vitro in 3D human iPS cell-engineered heart tissue. Nature communications. 8, 1078 (2017).

30 Smeets, J. L., Allessie, M. A., Lammers, W. J., Bonke, F. I. & Hollen, J. The wavelength of the cardiac impulse and reentrant arrhythmias in isolated rabbit atrium. The role of heart rate, autonomic transmitters, temperature, and potassium. Circulation research. 58, 96–108 (1986).

31 Izumi-Nakaseko, H. et al. Characterization of human iPS cell-derived cardiomyocyte sheets as a model to detect drug-induced conduction disturbance. The Journal of toxicological sciences. 42, 183–192 (2017).

32 Rensma, P. L., Allessie, M. A., Lammers, W. J., Bonke, F. I. & Schalij, M. J. Length of excitation wave and susceptibility to reentrant atrial arrhythmias in normal conscious dogs. Circulation research. 62, 395–410 (1988).

33 Courtemanche, M., Keener, J. P. & Glass, L. A delay equation representation of pulse circulation on a ring in excitable media. SIAM Journal on Applied Mathematics. 56, 119–142 (1996).

34 Courtemanche, M., Glass, L. & Keener, J. P. Instabilities of a propagating pulse in a ring of excitable media. Physical Review Letters. 70, 2182 (1993).

35 Shaw, R. M. & Rudy, Y. Ionic mechanisms of propagation in cardiac tissue: roles of the sodium and L-type calcium currents during reduced excitability and decreased gap junction coupling. Circulation research. 81, 727–741 (1997).

36 Beauchamp, P. et al. Electrical propagation in synthetic ventricular myocyte strands from germline connexin43 knockout mice. Circulation research. 95, 170–178 (2004).

37 Arduini, D., Rizzo, G. & Romanini, C. Fetal cardiac function. (Parthenon Pub. Group, 1995).

38 Groenendyk, J., Agellon, L. B. & Michalak, M. Coping with endoplasmic reticulum stress in the cardiovascular system. Annu. Rev. Physiol. 75, 49–67 (2013).

39 van der Velden, J., de Jong, J. W., Owen, V., Burton, P. & Stienen, G. Effect of protein kinase A on calcium sensitivity of force and its sarcomere length dependence in human cardiomyocytes. Cardiovascular research. 46, 487–495 (2000).

40 Asnes, C. F., Marquez, J. P., Elson, E. L. & Wakatsuki, T. Reconstitution of the Frank-Starling mechanism in engineered heart tissues. Biophys. J. 91, 1800–1810 (2006).

41 Stoppel, W. L., Kaplan, D. L. & Black Iii, L. D. Electrical and mechanical stimulation of cardiac cells and tissue constructs. Advanced Drug Delivery Reviews. 96, 135–155 (2016).

42 Lindsley, D. B. Heart and brain potentials of human fetuses in utero. The American Journal of Psychology. 55, 412–416 (1942).

43 DuBose, T. J., Cunyus, J. A. & Johnson, L. F. Embryonic heart rate and age. Journal of Diagnostic Medical Sonography. 6, 151–157 (1990).

44 Zhang, D. et al. Tissue-engineered cardiac patch for advanced functional maturation of human ESC-derived cardiomyocytes. Biomaterials. 34, 5813–5820 (2013).

45 Gwyther, T. A. et al. Engineered vascular tissue fabricated from aggregated smooth muscle cells. Cells Tissues Organs. 194, 13–24 (2011).

46 Boudou, T. et al. A microfabricated platform to measure and manipulate the mechanics of engineered cardiac microtissues. Tissue Engineering Part A. 18, 910–919 (2011).

47 Agarwal, A., Goss, J. A., Cho, A., McCain, M. L. & Parker, K. K. Microfluidic heart on a chip for higher throughput pharmacological studies. Lab on a Chip. 13, 3599–3608 (2013).

48 Marquez, J. P. et al. High-throughput measurements of hydrogel tissue construct mechanics. Tissue Engineering Part C: Methods. 15, 181–190 (2009).

49 Laflamme, M. A. & Murry, C. E. Heart regeneration. Nature. 473, 326–335 (2011).

50 Sato, T. et al. Effect of electrical stimulation on IGF-1 transcription by L-type calcium channels in cultured retinal Müller cells. Japanese journal of ophthalmology. 52, 217–223 (2008).

51 Ito, A. et al. Induction of functional tissue-engineered skeletal muscle constructs by defined electrical stimulation. Scientific reports. 4, 4781 (2014).

52 Kamioka, H., Maeda, E., Jimbo, Y., Robinson, H. P. & Kawana, A. Spontaneous periodic synchronized bursting during formation of mature patterns of connections in cortical cultures. Neurosci. Lett. 206, 109–112 (1996).

53 Huang, Y. J., Wu, H. C., Tai, N. H. & Wang, T. W. Carbon nanotube rope with electrical stimulation promotes the differentiation and maturity of neural stem cells. Small. 8, 2869–2877 (2012).

54 Shiba, Y. et al. Human ES-cell-derived cardiomyocytes electrically couple and suppress arrhythmias in injured hearts. Nature. 489, 322–325 (2012).

55 Minami, I. et al. A small molecule that promotes cardiac differentiation of human pluripotent stem cells under defined, cytokine-and xeno-free conditions. Cell Rep. 2, 1448–1460 (2012).

56 Minami, I. et al. A small molecule that promotes cardiac differentiation of human pluripotent stem cells under defined, cytokine-and xeno-free conditions. Cell reports. 2, 1448–1460 (2012).

57 Isomura, A., Hörning, M., Agladze, K. & Yoshikawa, K. Eliminating spiral waves pinned to an anatomical obstacle in cardiac myocytes by high-frequency stimuli. Physical Review E. 78, 066216 (2008).

58 Tripathi, S. et al. Meta-and orthogonal integration of influenza “OMICs” data defines a role for UBR4 in virus budding. Cell host & microbe. 18, 723–735 (2015).

## Reference

1. Priebe, L. & Beuckelmann, D. J. Simulation study of cellular electric properties in heart failure. Circ Res 82, 1206–1223 (1998).

2. Panfilov, a V. a. of Human Ventricular Cells. 281, 2296–2308 (2002).

3. Verkerk, A. O., Van Borren, M. M. G. J. & Wilders, R. Calcium transient and sodium-calcium exchange current in human versus rabbit sinoatrial node pacemaker cells. Sci. World J. 2013, (2013).

4. Karakikes, I., Ameen, M., Termglinchan, V. & Wu, J. C. Human Induced Pluripotent Stem Cell-Derived Cardiomyocytes: Insights into Molecular, Cellular, and Functional Phenotypes. Circ. Res. 117, 80–88 (2015).

5. Shaw, R. M. & Rudy, Y. Ionic mechanisms of propagation in cardiac tissue. Roles of the sodium and L-type calcium currents during reduced excitability and decreased gap junction coupling. Circ. Res. 81, 727–741 (1997).

